# PAYOFF-BIASED SOCIAL LEARNING UNDERLIES THE DIFFUSION OF NOVEL EXTRACTIVE FORAGING TRADITIONS IN A WILD PRIMATE

**DOI:** 10.1101/110221

**Authors:** Brendan J Barrett, Richard L McElreath, Susan E Perry

**Affiliations:** University of California, Davis; Animal Behavior Graduate Group; University of California, Davis; Department of Anthropology; Max Planck Institute for Evolutionary Anthropology; University of California, Los Angeles; Department of Anthropology; University of California, Los Angeles; Center for Behavior, Evolution, and Culture

**Keywords:** payoff-bias, social learning, behavioural traditions, *Cebus*, cultural transmission, extractive foraging

## Abstract

The type and variety of learning strategies used by individuals to acquire behaviours in the wild are poorly understood, despite the presence of behavioural traditions in diverse taxa. Social learning strategies such as conformity can be broadly adaptive, but may also retard the spread of adaptive innovations. Strategies like payoff-biased learning, in contrast, are effective at diffusing new behaviour but may perform poorly when adaptive behaviour is common. We present a field experiment in a wild primate, *Cebus capucinus*, that introduced a novel food item and documented the innovation and diffusion of successful extraction techniques. We develop a multilevel, Bayesian statistical analysis that allows us to quantify individual-level evidence for different social and individual learning strategies. We find that payoff-biased social learning and age-biased social learning are primarily responsible for the diffusion of new techniques. We find no evidence of conformity; instead rare techniques receive slightly increased attention. We also find substantial and important variation in individual learning strategies that is patterned by age, with younger individuals being more influenced by both social information and their own individual experience. The aggregate cultural dynamics in turn depend upon the variation in learning strategies and the age structure of the wild population.

## 1. Introduction

The existence of culture or behavioural traditions [1] in non-human animals has been a topic of intrigue to evolutionary biologists and ethologists for centuries [2–4]. Recently, research interest in animal cultures has soared, partially driven by findings from long-term cross-site collaborations within primatology [5–7] and cetaceology [8;9] in the early 21st century. As the diversity of taxa in which social learning is studied grows, it appears that traditions might be more widespread and ecologically meaningful than was previously appreciated.

As evidence accumulates, the study of cultural mechanisms has shifted focus from asking “can animals learn socially?” to “how and under what conditions do animals learn socially?” The ecological drivers that favour social learning are theoretically well explored [10]. The mechanistic details and evolutionary and ecological consequences of social learning are less well understood. From an individual’s perspective, it may be difficult to know whom or exactly what to copy. To cope with these difficulties, organisms use heuristics and strategies [10–12] to minimize the costs and increase the efficiency of social learning. Variation in learning strategy, whether between individuals or over the life course, may also be important [13–15].

Different strategies have different advantages. Two families of social learning strategies that have received both theoretical and empirical attention are conformity and payoff-bias [10;16;17]. Conformist transmission, or positive frequency dependence, can be adaptive especially in spatially heterogeneous environments [10;18;19]. However, unless it is combined with other, flexible strategies, conformity may prohibit more adaptive behaviours from spreading [18;20] or cause population collapse [21]. In contrast to conformity, payoff-biased social learning is very effective at spreading novel adaptations. Payoff-biased social learning attends to behaviour that is associated with higher payoffs and presumably increase fitness. However, it can be outperformed by conformity, once adaptive behaviour is common [22].

There is empirical evidence for both conformist and payoff-biased social learning in humans [17]. In other animals, conformity [23;24] has been studied more extensively than payoff-bias. To our knowledge, no non-human study has directly compared the explanatory power of conformity and payoff-biased social learning.

Here we report results from a field experiment with white-faced capuchin monkeys (*Cebus capucinus*) that is capable of distinguishing conformist and payoff-biased social learning. Capuchins are an excellent study system for understanding social learning and traditions. They are tolerant of foraging in proximity with conspecifics [25], independently evolved many brain correlates associated with intelligence [26;27] and display the largest recorded repertoire of candidate behavioural traditions of any platyrrhine: social conventions [7], interspecific interactions [28] and extractive foraging techniques [29–32]. Their reliance on social learning, frequency of innovation, and complexity of social interactions exemplifies what is predicted for long-lived animals with a slow life history strategy [33]. We investigated the innovation and transmission of extractive foraging techniques used to access the protected seeds of the *Sterculia apetala* fruit. This fruit occurs sporadically over the range of *C. capucinus*. Only some groups are experienced with it. By introducing the fruit to a naive group in controlled settings, we observed the rise and spread of new foraging traditions. We then inferred which social learning strategies best predict individual behaviour and how they influence the origins and maintenance of traditions.

The statistical analysis employs a multilevel (aka hierarchical or varying effects) dynamic learning model, of the form developed by [17], and inference is based upon samples from the full posterior distribution, using Hamiltonian Monte Carlo [34]. This model allows estimation of unique social and individual learning strategies for each individual in the sample. The analysis utilizes dynamic social network data which were available during each field experimental session. It also permits examination of the relationship between any individual state (i.e. age, rank) and learning strategy. The multilevel approach makes it possible to apply these models to field data that lack precise balance and repeatedly sample individuals. We provide all code needed to replicate our results and to apply this same approach to any group time series of behaviour.

We document that the capuchins innovated a number of successful techniques. However, these techniques vary in their physical and time requirements. The statistical analysis suggests that payoff-biased social learning was responsible for this spread of the quickest, most successful techniques through the group. We find no evidence of conformity, but do find evidence of weak anti-conformity—rare techniques attracted more attention. We also find evidence of an age bias in social learning, in which older individuals were more likely to be copied. Individuals varied in how they made use of social cues and individual experience, and age was a strong predictor. Our results comprise the first application of multilevel, dynamic social learning models to a study of wild primates and suggest that payoffs to behaviour can have important and different influences on social and individual learning. Methodologically, the approach we have developed is flexible, practical, and allows for a stronger connection between theoretical models of learning and the statistical models used to analyse data.

## 2. Study Design

### 2.1. Study system

This study was conducted between 2013 and 2015 on a group of habituated white-faced capuchin monkeys in and near Reserva Biologica Lomas Barbudal (RBLB) in northwest Costa Rica, during the months of December–February. See electronic supplemental material and [35;36] for additional information about field site.

Capuchins heavily rely on extractive foraging to exploit difficult to access resources; this makes them an excellent comparative study system for understanding the evolution of extractive foraging in humans [26]. In neotropical dry forests, capuchins increase their reliance on extractive foraging during seasonal transitions when resources are limited. Capuchins receive more close, directed attention from conspecifics when they are foraging on large, structurally protected foods [37]. Many of the techniques required to access protected foods are candidate behavioural traditions [29].

Panamà fruits, *Sterculia apetala*, are a dietary staple of capuchins at RBLB; they comprise 8% of the diet of most groups in the early dry season [37]. The fruits are *empanada* shaped and the fatty, protein rich seeds within are protected by a hardened outer husk and stinging hairs [38]. Instead of waiting for fruits to dehisce, capuchins will open closed fruits and work around their the structural defences, thus reducing competition with other organisms. Panamà fruits require multiple steps to effectively open, process, and consume, and panamà foraging generates the second highest level of close-range observation from conspecifics at RBLB [37]. Panamà processing techniques are also observed to vary between groups at RBLB and other field sites in the area [29], suggesting they are socially-learned traditions. Wild capuchins without prior exposure to panamà fruits can’t initially open them [38], suggesting that personal experience and/or social influence are important.

Panamà processing techniques differ in efficiency, measured by the average time it takes to open a fruit. Techniques also differ in efficacy, both in their probability of being successful and due to costs incurred by encountering stinging hairs. This contrasts with other extractive foraging traditions that show no difference in efficiency or efficacy [30].

The focal group of this study, Flakes group (N=25), fissioned from the original study group in 2003. They migrated to a previously unoccupied patch of secondary agricultural and cattle ranching land characterized by riparian forest, pasture and neotropical oak woodland, where panamà trees are almost non-existent as they typically grow in evergreen, primary forests. Group scan data collected on foraging capuchins at RBLB from 2003–2011 show that Flakes was never observed foraging panamà, whereas other groups spent up to 1.21% of their annual foraging time eating panamà (Table S1). Two trees were found in the territory during phenological surveys, but are at the periphery, have small crowns, and are in areas of the habitat shared with other capuchin groups. When this study was designed, veterans of the field site had no recollection of observing Flakes foraging for panamà. Observations of two natal Flakes adult males (old enough to be expert panamà foragers in any other group) found outside of their territory migrating, suggest that they had little or no experience with panamà fruits.

Five adults in the group (two females, three males) grew up in different natal groups whose territories contained large numbers of panamà trees and whose groups exhibited higher rates of panamà foraging. For two migrant males from non-study groups, it is unknown if they previously learned to process panamà, but this seems likely as evidenced by their skill. These individuals acted as models for different behaviours, as they differed in the primary panamà processing techniques they presumably acquired in their natal groups. By providing panamà fruits to both naöive/inexperienced juveniles and to knowledgeable adult demonstrators who differ in processing techniques, we collected fine-grained data showing how inexperienced capuchins learn a natural behaviour.

### 2.2. Data Collection

We collected panamà fruits from areas near RBLB for our experiment. Fruits were placed on a 25 cm diameter wooden platform which provided visual contrast of the fruits against the ground as fruits blended with the leaf litter, and so the capuchins had some sort of naturalistic spatial cue to associate with panamà fruits. Two fruits were placed on 1-2 platforms in each experimental bout. This permitted 1-4 capuchins to forage at a given time, and 2 fruits per platform was the maximum number on which a single human observer could reliably collect data.

We placed multiple fruits for two reasons. First, when individuals are naturally foraging for panamà, they choose from multiple available fruits in a tree. Second, we wanted to see whom they bias their attention toward when given a choice of multiple potential demonstrators. While many learning experiments have one potential demonstrator to learn from in a foraging bout or assume that everyone observes that demonstrator, we believe that allowing them to choose a potential learning model is more representative of how wild animals learn.

Fruits were placed on platforms under a poncho to obscure the monkey’s view of us handling fruits. As ponchos were worn regularly when not experimenting, monkeys were unlikely to associate their presence with panamà platforms. When monkeys were not looking, we uncovered the fruits and walked to an observation area away from the platform so the monkeys could forage unimpeded. On digital audio recorders, we recorded if or when individuals saw, handled, processed, opened, ingested seeds from, and dropped each fruit. We verbally described how they were processing each fruit using an ethogram of techniques and which audience members observed them. Further information about fruit collection, data collection, and observer training can be found in the electronic supplemental material text and video, in addition to descriptions (Table S2) and video of panamà processing techniques.

## 3. Statistical Analyses

We analysed these data using multilevel experience-weighted attraction (EWA) models [39;40]. EWA models are a family of models that link individual learning rules and social information use to population-level dynamics by fitting existing mathematical models of learning as statistical models [16;17;41].

### 3.1. Social learning strategies

Our main focus is the contrast between two well-studied types of social learning, conformity and payoff-bias. However, we also investigate other plausible strategies. We quickly describe the background of these strategies and how the modelling framework incorporates them.

#### Payoff-biased learning

Copying the behaviour with the highest observable payoff is a useful social learning strategy [22;42]. In a foraging context, selectively copying rate-maximizing behaviour can increase the efficiency of diet and resource acquisition. Guppies choose food patches with higher return rates [43] while wild tufted capuchins bias attention toward the most efficient tool-users [44]. Cues of payoff may be noisy, however, and different individuals may require different techniques.

#### Model-biased learning

Sometimes evaluating the content of a behaviour is costly or impossible. In these circumstances, it may be an adaptive heuristic to bias attention toward particular demonstrators or “models,” who display cues (i.e. rank, health, fertility) that are likely to be correlated with adaptive behaviour.

Prestige-biased learning is a popular example of model bias in humans [45]. While animals may lack the concept of prestige, they have analogues. Captive chimpanzees have been found to be more likely to copy dominant individuals [41;46], while vervets copy same-sex high-ranking individuals [47].

Copying the behaviour of one’s parents is another option. If a parent can survive and successfully reproduce, its offspring’s existence serves as a cue that her parents are successful [48]. *Luehea* processing techniques of capuchins at RBLB were predicted by both the technique their mother used and the technique they saw performed most often [30]. Kin-biased learning has been found in carnivores [49–51], but it is unclear whether this is due to cognition or is a consequence of family-unit social systems.

Copying similar individuals can be adaptive. Where individuals differ in strength, size, or cognitive ability, it might be beneficial for learners to copy those who are most similar to them. Sex-biased learning has been found in several primate species [30;47].

#### Frequency-dependent learning

Frequency-dependent social learning occurs when frequency among demonstrators or frequency of demonstration influences adoption. It includes negative and positive frequency-dependence. Negative frequency dependence, or anti-conformity, is preferentially copying rare behaviour. It may be a form of neophilia. Positive frequency dependence, known also as conformity or majority-rule, is preferentially copying the most common behaviour. Conformity can lead to the fixation and maintain the stability of a cultural trait [10;18]. Experiments in many captive [20;52–55] and some wild [23;24] animals have found evidence of conformist learning.

## 3.2. Model design

An EWA model comprises two parts: a set of expressions that specify how individuals accumulate experience and a second set of expressions that specify the probability of each option being chosen. Accumulated experience is represented by *attraction* scores, *A*_*ij,t*_, unique to each behaviour *i*, individual *j*, and time *t*. A common formulation is to update *A*_*ij,t*_ with an observed payoff *π*_*ij,t*_:

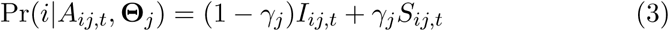

The parameter *ϕ*_*j*_ controls the importance of recent payoffs in influencing attraction scores. This parameter is unique to individual *j*, and so can vary by age or any other feature.

To turn these attraction scores into behavioural choice, some function that defines a probability for each possible choice is needed. The conventional choice is a standard multinomial logistic, or *soft-max*, choice rule:

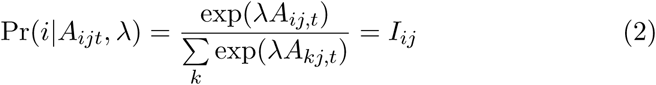

The parameter *λ* controls how strongly differences in attraction influence choice. When *λ* is very large, the choice with the largest attraction score is nearly always selected. When *λ* = 0, choice is random with respect to attraction score. Individuals were assigned a payoff of zero, *π*_*ij,t*_ = 0, if they failed to open a panamà fruit. If they were successful, payoff was the inverse-log amount of time it took to open the fruit, *π*_*ij,t*_ = log(*T*_open_)^−1^. For the observed times *T*_open_, this ensures that payoffs decline as *T*_open_ increases, but with the steepest declines early on.

Following previous work, social learning may influence choice directly and distinctly from individual learning. Let *S*_*ij*_ = *S*(*i*|**Θ**_*j*_) be the probability an individual *j* chooses behaviour *i* on the basis of a set of social cues and parameters **Θ**_*j*_. Realized choice is given by:

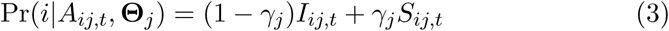

where *γ*_*j*_ is the weight, between 0 and 1, assigned to social cues. Under this formulation, social cues influence choice directly but attraction scores indirectly, only via the payoffs choice exposes an individual to.

We incorporate social cues into the term *S*_*ij,t*_ by use of a multinomial probability expression with a log-linear component *B*_*ij,t*_ that is an additive combination of cue frequencies. Specifically, the probability of each option *i*, as a function only of social cues, is:

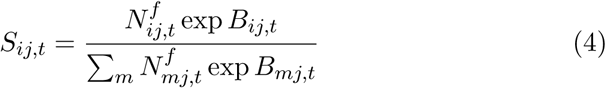

This is easiest to understand in pieces. The *N*_*ij,t*_ variables are the observed frequencies of each technique *i* at time *t* by individual *j*. The exponent *f* controls the amount and type of frequency dependence. When *f* = 1, social learning is unbiased by frequency and techniques influence choice in proportion to their occurrence. When *f >* 1, social learning is conformist. Other social cues, like payoff, are incorporated via the *B*_*ij,t*_ term:

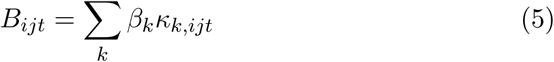

This is the sum of the products of the influence parameters *β*_*k*_ and the cue values *ϰ*_*k,ijt*_. We consider five cues:

(1) Payoff. *ϰ* = log(*t*_open_)^−1^ or, for failure, *ϰ* = 0.
(2) Demonstrator rank. *ϰ* = 1 for alpha rank, 0 otherwise.
(3) Matrilineal kinship. *ϰ* = 1 for matrilineal kin, 0 otherwise.
(4) Age similarity. *ϰ* is defined as the inverse absolute age difference: (1 + |age_demonstrator_ -age_observer_*|*)^−1^.
(5) Age bias. *ϰ* = age_demonstrator_.

The final components needed are a way to make the individual-level parameters depend upon individual state and a way to define the window of attention for social cues at each time *t*. The parameters *γ*_*j*_ and *ϕ*_*j*_ control an individual *j*’s use of social cues and rate of attraction updating, respectively. We model these parameters as logistic transforms of a linear combination of predictors. For example, the rate of updating *ϕ*_*j*_ for an individual *j* is defined as:

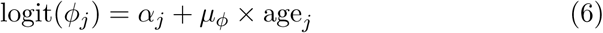

where *α*_*j*_ is a varying intercept per individual and *μ*_*ϕ*_ is the average influence of age on the log-odds of the updating rate. Social information available at each time step in the model was a moving window of the previous 14 days of observed foraging bouts. This allows new social information to be used, while old information is discarded. We tested the sensitivity of the time window used to calculate social cues and found our results were robust to variations in window width (7, 14, 21, 28 days) (Table S4). Attempts to parametrise window width fit poorly. To fit the model, we defined a global model incorporating all cues, using both parameter regularization and model comparison with sub-models to account for overfitting. Overall nine models were fit representing nine learning strategies. Models were fit using the Hamiltonian Monte Carlo engine Stan, version 2.14.1 [34], in R version 3.3.2 [56]. We compared models using WAIC [57]. To check our approach, we simulated the hypothesized data generating process and payoff structure and recovered data-generating values from our simulated data. We chose conservative, weakly informative priors for our estimated parameters. This made our models sceptical of large effects and helped ensure convergence.

## 4. Results: Innovation and Diffusion of Techniques

Of the 25 individuals in the group, 23 tried to process panamà and 21 were successful at least once over 75 experimental days. We observed 7 types of predominant fruit processing techniques on 1441 fruits, which varied in time required and the proportion of successful attempts (Table S2). Mean (median) duration ranged from 50 (29) to 330 (210) seconds. Proportion of successful attempts ranged from 0.38 to 0.89.

**Figure 1.**
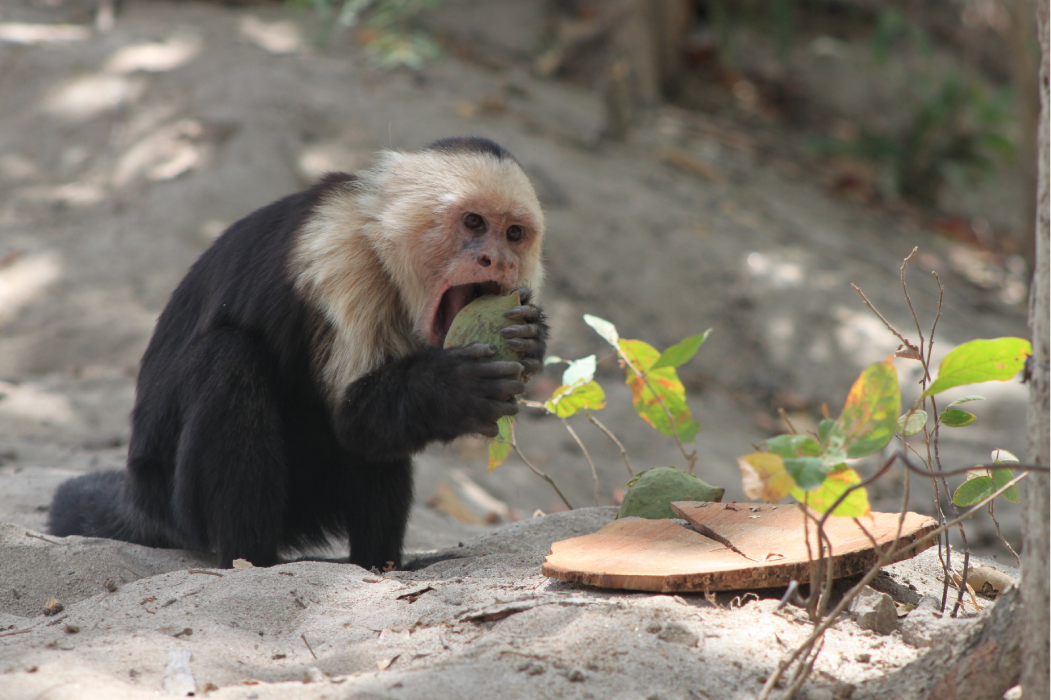
Adult male NP exhibits the canine seam technique.

The technique frequencies changed over time, in the group and in most individuals (figures 2,S3,S4). The most efficient technique, canine seam, went from non-existent in the group to the most common technique. It was introduced by an immigrant adult male (NP). Two knowledgeable adults, an adult female (ME) and the alpha male (QJ), switched to the canine seam technique. All others born after 2009 tried it at least once (figure S4). However, canine seam never reached fixation in the population.

### 4.1. Results of EWA models

There was overwhelming support for some mix of individual and social learning over individual learning alone (Table S3). The highest ranked model was the global model containing all strategies and age effects on learning parameters, which received 94% of the total model weight. We focus on this model, as it is both highest ranking and its parameter values agree with the weights assigned in the overall model set.

Marginal posterior distributions of each parameter are displayed in Table 1 and visualized in figure S1. Note that the marginal posterior distribution of each parameter cannot be directly interpreted as the importance of each factor in the total diffusion of behaviour. The weight of social information (*γ*), for example, can be relatively small at each instantaneous choice but still be decisive in determining which behaviour spreads, because individual discovery rates may be even smaller. As each individual’s behaviour is unique to their observed social information, personal experience, and estimated individual-level parameters, we encourage readers to view marginal predictions with visualizations of implied individual behaviour, using posterior predictive distributions in the electronic supplementary material (figure S3).

**Figure 2.**
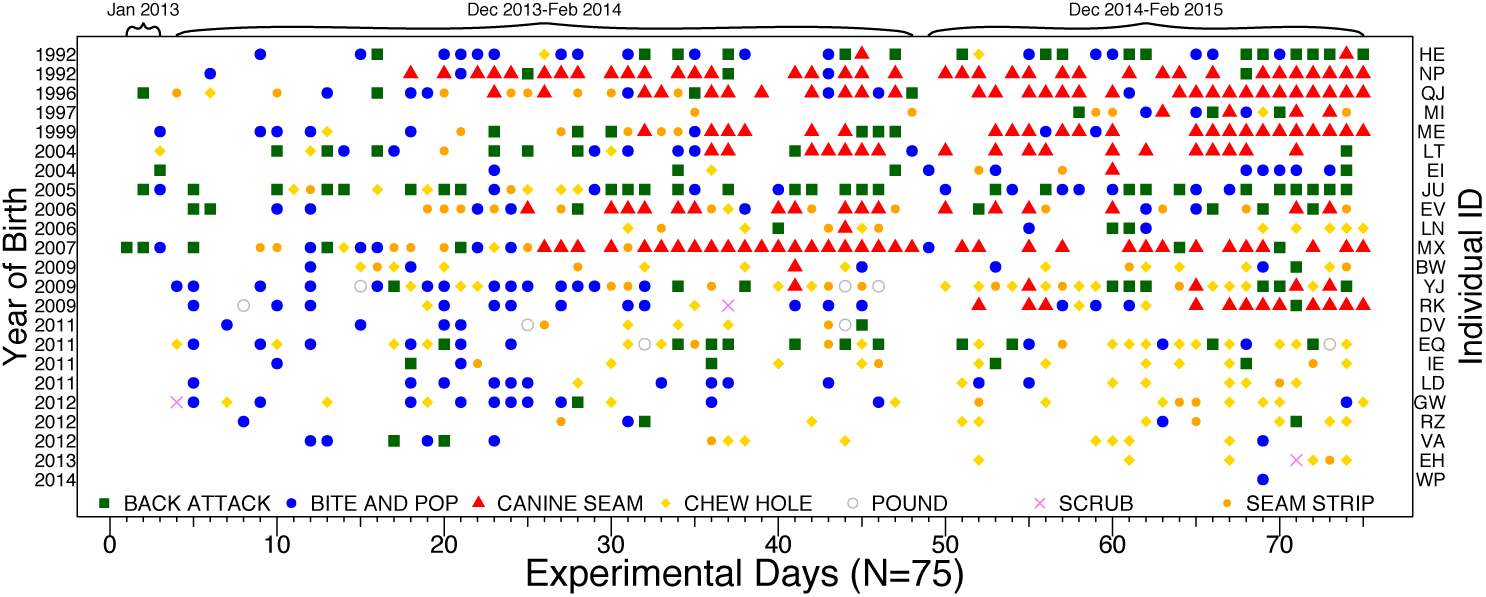
Techniques observed during experiment. Rows are unique individuals, from oldest (top) to youngest (bottom). X-axis is sequential order of experimental days. Each color/shape represents most common technique used by an individual on that day, no point indicates days of no processing. The most successful technique indicated by red triangles (canine seam) diffused to older members of the population. Younger individuals did not use canine seam.

#### Influence of conformity and payoff-bias (*f* and *β*_pay_)

The raw marginal conformist exponent is below 1 on average, indicating mild anticonformity—a bias toward copying rare behaviours. The marginal payoff-bias coefficient is strongly positive, indicating attraction to high-payoff actions. Figure 3 visualizes the individual social learning function *S*_*ijt*_ (Expression 4) implied when only conformity and payoff-bias are present. The horizontal axis is the observed frequency of a higher payoff option among demonstrators. The vertical axis is the probability an individual chooses the higher payoff option. Each curve in the figure represents the posterior mean for an individual. The diagonal dashed line represents unbiased social learning. All individuals are strongly biased by payoff, resulting in a preference for the high-payoff option over most of the range of the horizontal axis. But most individuals also display weak anti-conformity, resulting in a preference for the rarer, low-payoff option in the upper right corner.

#### Weight of past experience (*ϕ*)

On average, capuchins more heavily favor previous experiences over new ones (*ϕ* = 0.15; [0.11, 0.20] 89% credible interval), Table 1). However, there is considerable individual variation in attraction to new experience (*σ*_*individual*_ = 0.66) ranging from 0.08 to 0.36, which was negatively predicted by age (*μ*_*age*_ = –0.11; 89% CI [–0.16, *–*0.06]; figure 4a). This suggests that older individuals are more canalized than younger individuals.

**Table 1.**
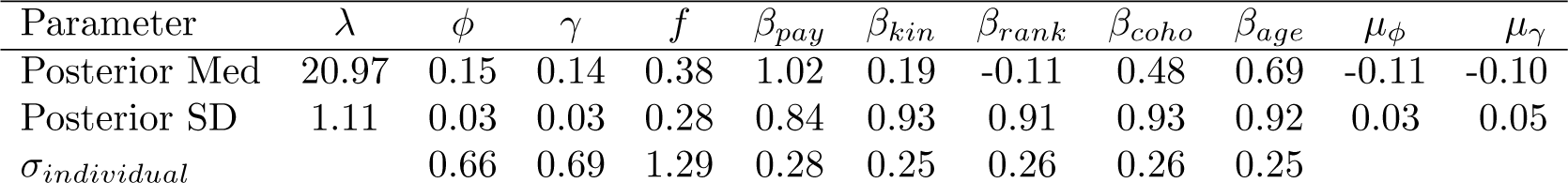
Posterior medians and standard deviations from the global model. Estimates of *σ*_*individual*_ are the standard deviations of varying effects for that parameter across individuals. Posteriors visualized in figures S1 and S2.

| Parameter | $\lambda$ | $\phi$ | $\gamma$ | $f$ | $\beta_{pay}$ | $\beta_{kin}$ | $\beta_{rank}$ | $\beta_{coho}$ | $\beta_{age}$ | $\mu_\phi$ | $\mu_\gamma$ |
| --- | --- | --- | --- | --- | --- | --- | --- | --- | --- | --- | --- |
| Posterior Med | 20.97 | 0.15 | 0.14 | 0.38 | 1.02 | 0.19 | -0.11 | 0.48 | 0.69 | -0.11 | -0.10 |
| Posterior SD | 1.11 | 0.03 | 0.03 | 0.28 | 0.84 | 0.93 | 0.91 | 0.93 | 0.92 | 0.03 | 0.05 |
| $\sigma_{individual}$ | | 0.66 | 0.69 | 1.29 | 0.28 | 0.25 | 0.26 | 0.26 | 0.25 | | |

#### Weight of social information (*γ*)

*γ* estimates for individuals varied considerably, ranging from 0.07-0.39 (*σ*_*individual*_ = 0.66). *γ* was also negatively related to age (*μ*_*age*_ = –0.10; 89% CI [–0.18, *–*0.03]; Fig. 4b). This suggests that younger individuals rely more on social cues.

#### Age bias (*β*_*age*_)

Age bias contributed notably to social learning in our global model (*β*_*age*_ = 0.69; 89% CI [–0.79, 2.14]; Table 1), suggesting that all capuchins were more likely to copy older demonstrators.

#### Age similarity, kin, and rank biases

None of age similarity, matrilineal kin, or rank biases presented a strong or consistent effect (coho, kin, and rank in Table 1). While these strategies may have influenced some individuals and decisions, there is little evidence of general importance for these cues.

## 5. Discussion

We set out to examine the roles of conformist and payoff-biased social learning among wild capuchin monkeys during the diffusion of novel food processing techniques. We find no evidence of conformity, defined as positive frequency dependence. We do however find strong evidence of payoff-biased learning.

Little work has examined whether animals use payoff-biased social learning. We do not know how common such strategies are in nature. It is common to experimentally examine payoff-equivalent options, shedding no light on payoff-bias. The common exclusion approach to identifying animal culture accidentally excludes payoff-bias, by diagnosing ecologically correlated behavioural differences as non-cultural [5]. This may result in overlooking adaptive socially-learned behaviour. If payoff-bias is common, this makes the problem of identifying animal traditions more subtle.

We also found evidence that other social cues, such as age, influence social learning. Age also modulated underlying learning parameters. In combination, these influences are sufficient to describe the diffusion and retention of successful foraging techniques within the group. In the remainder of the discussion, we elaborate on the findings and summarize some of the advantages and disadvantages of our approach.

**Figure 3.**
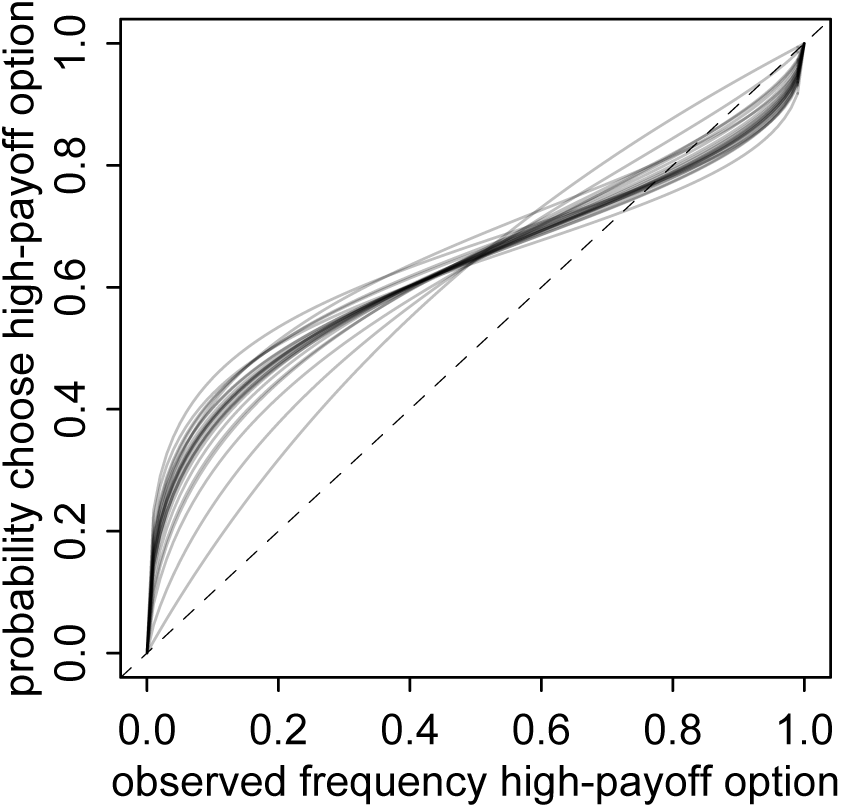
Posterior predictions of probabilities of choosing a socially observed option with payoff log(*t*_open_)^−1^ = 0.5, relative to an observed option that was not successfully opened.

**Figure 4.**
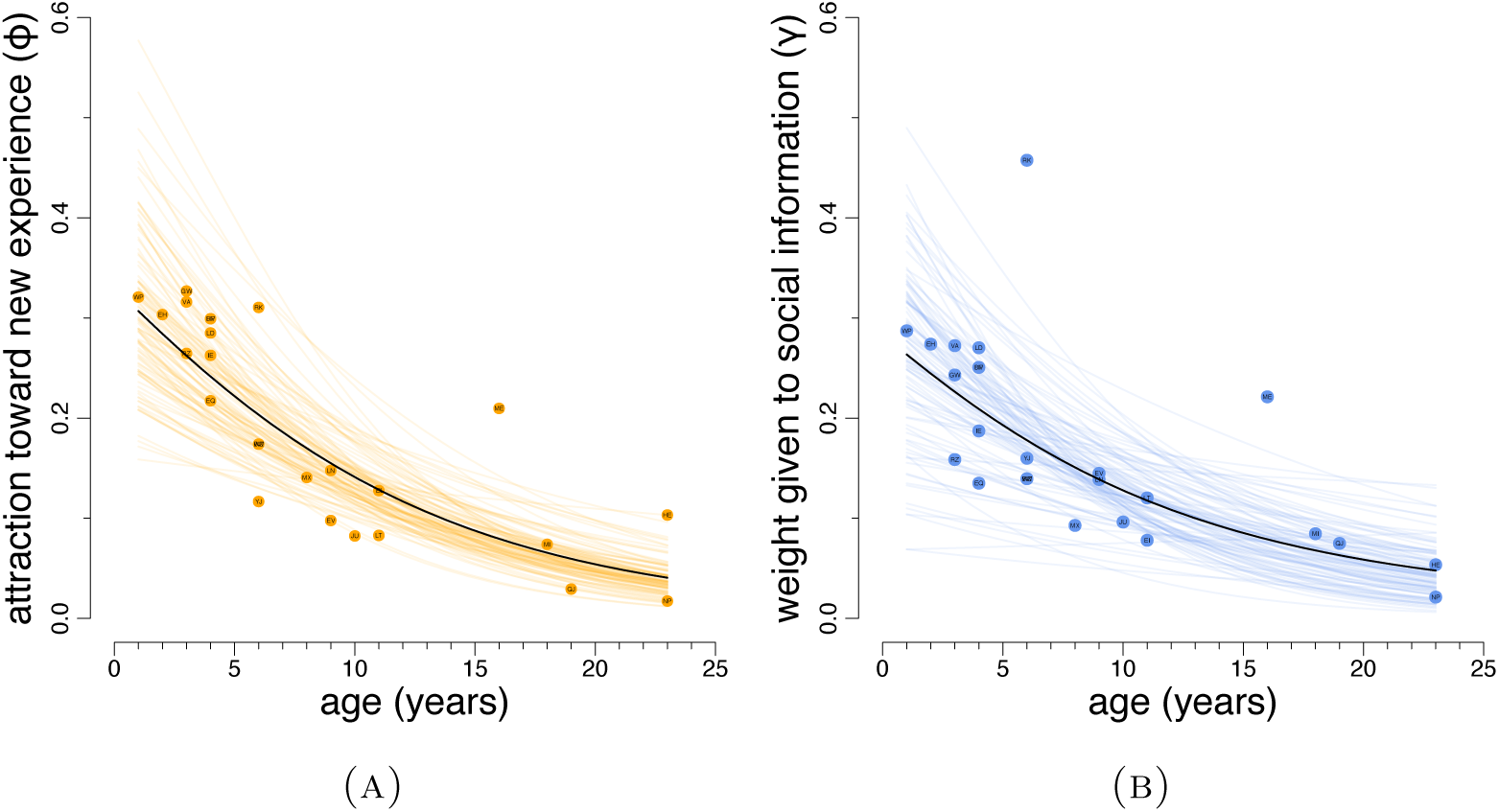
Relationships between age and (A) attraction to new experience (*ϕ*) and (B) influence of social information (*γ*). Black line represents posterior mean. Solid points are posterior means of individual varying effects. Lighter lines are 100 posterior samples.

### 5.1. Wild capuchins acquire extractive foraging techniques quickly via social learning

This study shows that one group of wild capuchin monkeys socially learn extractive foraging techniques from conspecifics and supports claims that food processing techniques are socially learned traditions. It has been challenging to find experimental evidence for social learning of object manipulation tasks in captive capuchins [26;58]. Better evidence for social learning might be found across a broader range of taxa if more ecologically valid behaviours are studied in the wild. This study also demonstrates that capuchins, like other animals [59], may be able to acquire new, efficient behaviour in a matter of days or weeks if knowledgeable models are available. This rapid pace of social transmission suggests that learning can act to rapidly facilitate behavioural responses to environmental change [12].

We found that payoff-biased learning and negative frequency dependence guided diffusion of panamà processing techniques in this group (Table 1). These strategies are consistent with the observation that the rarest and most efficient panamà processing technique, canine seam, eventually became the most common. This was the case for most, but not all, naive and knowledgeable adults and subadults born after 2009 (figure 2). Juveniles born before 2009 did not use the canine seam technique (figure S4 and 2), likely because their mouths were not sufficiently large and strong.

Payoff-bias had the largest effect on the probability of choosing a behaviour, while negative frequency dependence may have prevented it from ever reaching fixation. Experimental evidence of wild animals using payoff-biased learning has not been previously reported. Our finding of negative frequency-dependent learning suggests that capuchins bias their attention towards rare or novel behaviours—a type of neophilia.

While all adult individuals tried the canine seam technique, they typically settled on the technique(s) that was most successful for them. Individuals who settled on the canine seam technique also sporadically tried other behaviours (figure S4). This result is consistent with other research [60] suggesting that social learning guides exploration but personal experience strongly influences adoption.

While we found the strongest support for payoff-biased learning, our modelling suggests that animals use multiple social learning strategies simultaneously or that social biases and content biases might be equifinal. Age-biased learning also had support in the global model (Table 1). This might be due to older individuals’ increased likelihood of being efficient panamà processors compared to juveniles, but the preferences for some individuals (JU and LN) to copy the techniques of the adults they commonly associate with who did not use canine seam (HE and MI respectively) suggests otherwise.

Nevertheless, observational studies are always limited in their ability to distinguish some mechanisms from others. We believe that long-term field studies, field experiments, and controlled captive experiments all have important and complementary roles to play.

### 5.2. Age predicts individual variation in social and individual learning

Individual variation in social learning may have meaningful evolutionary and social implications, yet remains poorly studied [13]. We found that younger individuals more heavily relied on social learning than older individuals (figure 4b) and that older individuals were less likely to observe conspecifics (figure S5).

We also observed that older individuals were less likely to update information and had a greater attraction to previous experiences (figure 4a). This might be due to older individuals being less exploratory than younger individuals. One alternative explanation is that older individuals’ higher success rates at processing panamà provided them with higher quality personal information to discern between the efficiency of varied processing techniques (figure S4). This age structure in proclivity to learn socially suggests flexible learning strategies that change over development. Theory predicting and explaining such flexible variation waits to be constructed.

### 5.3. Statistical approach

Our analytical approach was designed around three important principles. First, it allows us to evaluate the possible influence of several different, theoretically plausible, social learning biases. Second, the framework combines social learning biases with a dynamic reinforcement model in which individuals remember and are influenced by past experience with different techniques. Third, the approach is multilevel, with each individual possessing its own parameters for relative use of each learning strategy. This allows us to evaluate heterogeneity and its contribution to population dynamics.

Our approach is distinct from looking for evidence of population-level learning dynamics consistent with the hypothesized learning strategy (i.e sinosoidal curves and conformity) [24;61]. In our approach any population-level patterns are consequences of inferred (and potentially different) strategies among individuals (visualized in figure S3); they are not themselves used to make inferences about learning.

Our approach is most similar to network-based diffusion analysis (NBDA) [62;63]. In principle, our framework and NBDA can be analogized, despite differences in the details of modelled strategies, because both are multinomial time series modelling frameworks that can be treated as both survival (time-to-event) or event history analyses. There are some notable differences in practice. Our approach differs from typically employed NBDA in that it: 1) uses a full dynamic time series for available social information rather than a static social network and 2) emphasizes modelling the entire behavioural sequence including and beyond the first putative instance of social transmission. There is no reason in principle that ordinary NBDA models could not make similar use of these data, and recent advances [59] utilize dynamic social networks.

It is important to note that successfully fitting these dynamic, multilevel models benefits from recent advances in Monte Carlo algorithms. We used an implementation of Hamiltonian Monte Carlo (NUTS2) provided by Stan [34]. Our global model contains 231 parameters and would prove very challenging for older algorithms like Gibbs Sampling. Hamiltonian Monte Carlo not only excels at high-dimension models, even with thousands of parameters, but it also provides greatly improved mixing diagnostics that allow us to have greater confidence in the correctness of the results, regardless of model complexity.

### 5.4. Implications for the origins and maintenance of traditions

This model suggests that payoff-biased learning can cause the spread of a tradition. However, social learning may increase within-group homogeneity, while individual learning may act to decrease it [51]. Our findings are consistent with this idea. Limited transfer of individuals in xenophobic species like *Cebus* is exceptionally important in maintaining group specific traditions for behaviours that differ in payoff. However, this likely acts concordant with transmission biases. Variation might also be maintained due to biases for copying particular subsets of individuals (e.g. a particular age-class or kin group) in a stable social system. Migration of new individuals with more efficient behaviours could seed a new tradition in the group, the diffusion of which may be due to payoff-biased learning.

### 5.5. Future Directions

We have noted that equifinality might exist between learning strategies. On average, older individuals were better at opening panamà fruit. Perhaps individuals are biasing learning toward older individuals and acquiring the efficient techniques indirectly instead of turning attention toward the content of the behaviour. While we think this is likely not the case based on the evidence considered in this study, it is a possibility in all learning studies. In many cases, where we are interested in predicting the population dynamics of learning in a given context, the exact social learning strategy might not matter if it has the same dynamics and leads to the same frequency in a population. Many learning strategies are likely equifinal under the right social conditions. However, the exact nature of the cognitive mechanisms of the learning strategies organisms employ, and the social factors which indirectly structure learning, become important when we wish to use social learning in applied contexts. Further theoretical and empirical explorations of social learning need to address that learning is a two stage process: one of assortment and one of information use.

An important aspect of learning that we have neglected is the endogeneity of social information. Our statistical models evaluated how individuals use information they observed. However, before individuals acquire social information, they make the decision to observe others. Future analyses will evaluate who individuals choose to bias attention toward when in the proximity of potential demonstrators to see how positive assortment due to social preferences, rank, or food sharing might structure opportunities for social learning and affect the establishment and maintenance of traditions.

Most models of social learning in the evolutionary anthropology and animal behaviour literature assume a randomly assorted population. However, non-random assortment occurs before information is acquired in a population, and it can drastically affect social learning and cultural dynamics. Sometimes this assortment may be an adaptive heuristic, such as deciding to bias attention. Other times it may be an indirect consequence of social behaviour, such as avoidance of a potentially dangerous demonstrator [15]. Asymmetrical age structure in a population may also make the behavioural variants in the population non-random when learning abilities are constrained by skill and developing cognition [64]. Social networks can also change drastically over development, opening up avenues for new possible learning strategies. Some learning strategies might be difficult to tease apart in small, non-diverse social systems. If a juveniles engage in kinbiased learning [65], but only interact with their kin group, how are we to discern kin-biased learning from linear imitation or conformity, and under what conditions does this distinction matter?

## 6. Authors’ Contributions

BB designed study, collected data, carried out analysis, and drafted manuscript; RM participated in analysis and helped draft manuscript; SP established field site, collected data, and helped draft manuscript.

## 7. Acknowledgements

Thanks to: M. Grote, M. Crofoot, and the UC Davis CE/HBE lab for useful feedback, A. Cobden, B. Davis, E. Seabright, M. White, M. Ziegler, C. Angyal, for assistance with data collection, W. Lammers, J.C Ordoñez Jimènez, I. Godoy, and K. Kajokaite, for assistance with field logistics, and MINAET and SINAC, Hacienda Pelon de la Bajura, Hacienda Brin D’Amor for permission to work on their land. Additional acknowledgements are in the electronic supplemental material. This research complied with Costa Rican law, and followed UCLA (ARC #1996-122 and 2005-084 plus renewals) and UC Davis (IACUC permit #17297) animal care and use protocols.

## 8. Research Funds

Funding to BB was provided by the American Society of Primatologists, the ARCS Foundation, and a NSF GRF (Grant No. 1650042). SP received funding from the Max Planck Institute for Evolutionary Anthropology, the National Science Foundation (grants No. SBR-0613226 and BCS-0848360), Leakey Foundation, National Geographic Society, and UCLA COR. Any opinions, findings, and conclusions or recommendations expressed in this material are those of the authors and do not necessarily reflect the views of the National Science Foundation.

## 9. Data Accessibility

Data and code for models, simulations, and graphs are available at https://github.com/bjbarrett/panama1.

